# N-3 polyunsaturated fatty acids promote astrocyte differentiation and neurotrophin production independent of cAMP in patient-derived neural stem cells

**DOI:** 10.1101/2020.01.22.916130

**Authors:** Jiang-Zhou Yu, Jennifer Wang, Steven D. Sheridan, Roy H. Perlis, Mark M. Rasenick

## Abstract

Evidence from epidemiological and laboratory studies, as well as randomized placebo-controlled trials, suggests supplementation with n-3 polyunsaturated fatty acids (PUFAs) may be efficacious for treatment of major depressive disorder (MDD). The mechanisms underlying n-3 PUFAs potential therapeutic properties remain unknown. There are suggestions in the literature that glial hypofunction is associated with depressive symptoms and that antidepressants may normalize glial function. In this study, iPSC-derived neuronal stem cell lines were generated from individuals with MDD. Astrocytes differentiated from patient-derived neuronal stem cells (iNSCs) were verified by GFAP. Cells were treated with eicosapentaenoic acid (EPA), docosahexaenoic acid (DHA) and stearic acid (SA). During astrocyte differentiation, we found that n-3 PUFAs increased GFAP expression and GFAP positive cell formation. BDNF and GDNF production were increased in the astrocytes derived from patients subsequent to n-3 PUFA treatment. Stearic Acid (SA) treatment did not have this effect. CREB activity (phosphorylated CREB) was also increased by DHA and EPA but not by SA. Furthermore, when these astrocytes were treated with n-3 PUFAs, the cAMP antagonist, RP-cAMPs did not block n- 3 PUFA CREB activation. However, the CREB specific inhibitor (666-15) diminished BDNF and GDNF production induced by n-3 PUFA, suggesting CREB dependence. Together, these results suggested that n-3 PUFAs facilitate astrocyte differentiation, and may mimic effects of some antidepressants by increasing production of neurotrophic factors. The CREB-dependence and cAMP independence of this process suggests a manner in which n-3 PUFA could augment antidepressant effects. These data also suggest a role for astrocytes in both MDD and antidepressant action.

## Introduction

Major depressive disorder (MDD) is the most common psychiatric disorder, with almost one in six individuals experiencing at least one depressive episode at some point in their lifetime. It is currently the leading cause of disability worldwide [1, 2]. While effective treatments exist, about one third of patients treated with antidepressants do not reach symptomatic remission [3, 4]. While some of these non-remitters may respond to ketamine or other rescue strategies, additional therapeutic options are needed.

Brain regions and even cell-types contributing to MDD are not well established [4]. Evidence from clinical, preclinical and post-mortem studies suggests that dysfunction and degeneration of astrocytes may be one of potential candidates, and astrocytes may also represent a potential therapeutic target for MDD [5–8]. Antidepressants, including SSRIs and ketamine, increase brain-derived neurotrophic factor (BDNF) and glial cell-derived neurotrophic factor (GDNF) expression in primary cultured animal astrocytes [9–11]. Substantial evidence suggested that astrocytes might be the main resource of antidepressant-induced BDNF [8]. Furthermore, overexpression of BDNF in astrocytes leads to antidepressant-like activity in mice [12].

N-3 polyunsaturated fatty acids (n-3 PUFAs) are found in fish oil, and include EPA and DHA. Many studies have suggested antidepressant efficacy for omega-3 fatty acids (n-3 PUFA) in MDD [13–17], and n-3 PUFAs might evoke an antidepressant paradigm in culture cells and animal models [13]. The mechanisms of n-3 PUFA antidepressant action remain poorly documented, but include inhibition of the production of proinflammatory cytokines and maintenance of neuronal membrane stability and fluidity [13, 18]. Most efforts to characterize the mechanism of action of n-3 PUFAs have been done in animals and cell lines. Further, probing the mechanisms of n-3 PUFA anti-depressant effects, especially in human cells, would lead to better strategies for the use of n-3 PUFAs to improve outcome of MDD.

The restricted access to brain tissue and a variety of neuronal cell types in patients represents a major hurdle in exploring neuropsychiatric diseases mechanisms and developing therapeutic targets and approaches. Human-induced pluripotent stem cell (iPSC) technology offers another means of accessing neuropsychiatric patient cells by enabling the differentiation of patient iPSCs into relatively pure populations of neurons or glial cells, which may provide new avenues to study neuropsychiatric disorders and potential treatments [19]. The neuronal or glial cells resulting from patient-derived iPSC have been successfully used to investigate bipolar disorder, schizophrenia and MDD[20] [19, 21–23].

To explore the potential mechanisms underlying the beneficial effects of n-3 PUFAs in MDD, human astrocytes were obtained by differentiating MDD patient-derived iNSCs from iPSCs. N-3 PUFAs facilitated astrocyte generation from iNSCs during differentiation. The astrocytes also responded to n-3 PUFA with increased CREB activation and increased neurotrophin production. These were cAMP-independent, even though evidence suggested the importance of cAMP signaling in therapeutic effect of antidepressants[24]. This raises the possibility that, while n-3 PUFA have a cellular hallmark similar to antidepressants, they achieve this through a different pathway.

## Materials and methods

### Subject Recruitment

Two patient iPSC lines were drawn from a large biobank collected between 2010 and 2018 at Massachusetts General Hospital. Briefly, adult participants were recruited from outpatient psychiatry clinics and signed written, informed consent approved by the Partners HealthCare institutional review board. Diagnosis was confirmed by structured clinical interview by a trained physician rater [25]. Current and lifetime treatment history was collected using the Antidepressant Treatment Response Questionnaire[26] and the FAST[27], augmented with clinician review of electronic health records. Selective serotonin reuptake inhibitor sensitive (SSRI-sensitive) was defined as remission with one SSRI. SSRI-resistant was defined as lack of remission with two adequate SSRI trials. Participants underwent a 3 or 4mm dermal punch which was subsequently expanded into a fibroblast cell line. iPSC lines were generated from cultured fibroblast lines as previously described [20].

### Human iNSC culture and astrocyte differentiation

iNSCs from 1 SSRI-sensitive and 1 SSRI-resistant MDD patient were generated from iPSC lines cultured feeder free in E8 medium (Gibco) on Geltrex (Gibco) coated 6-well plates and passaged with Accutase (Sigma). iPSCs were cultured in E8 with 10uM Thiazovivin (Stemgent, Inc) following passaging. Prior to neural induction, iPSCs were purified using magnetic-activated cell sorting (MACS) with Tra1-60 microbeads (Miltenyi Biotec) on LS columns as described by vendor and cultured to 10-20% confluence. Neural induction was initiated using Neurobasal medium (Gibco) supplemented with 1X Neural Induction Supplement (Gibco). Induced cells were cultured to confluence in neural expansion medium (NEM; 50% Neurobasal, 50% Advanced DMEM/F-12, 1X Neural Induction Supplement; Gibco) and passaged using Accutase with 10uM Thiazovivin for at least 6 passages. Putative iNSCs were further purified by sequential MACS (PSA-NCAM^+^, CD271^-^, CD133^+^; Miltenyi Biotec) according to the manufacturer’s protocol.

Glial fibrillary acidic protein (GFAP)-expressing astrocytes were differentiated from iNSCs as reported with modifications [28, 29]. Briefly, iNSC were seeded in 6-well dishes coated with Geltrex at a seeding density of 5×10^5^/ml in NEM. On the second day, NEM was replaced by astrocyte differentiation medium (DMEM/F-12, 5ng/ml BMP2, 5ng/ml CNTF, and 10 ng/ml bFGF (PeproTech, Rock Hill, NJ)). Medium was exchanged every other day.

For testing the effect of n-3 PUFAs on astrocyte differentiation from iNSC, cells were seeded in NEM on Geltrex-coated 24-well plates or culture dishes. After 24 hours, the medium was changed to astrocyte differentiation medium containing vehicle (ethanol, 0.15%), docosahexaenoic acid (DHA, 50 μM), eicosapentaenoic acid (EPA, 50μM) or stearic acid (SA, 50 μM), prepared as previously reported [30]. After 5 days, cells were processed for qPCR, western blot assay or immunostaining. For studying the impact of n-3 PUFAs on neurotrophin production, astrocytes were differentiated for 30 days from iNSCs and then treated with vehicle, DHA, EPA or SA (same concentration as above described) for 3 days in 6–well plates and then processed to designed experiments.

### Immunocytochemistry

After fixation with cold 100% methanol (–20 °C) for 10 min, cells were incubated with blocking buffer (PBS, 5% goat serum, 0.3% Triton X-100) for 60 min, then incubated in 1:100 dilution of anti-PAX6 (Cell Signaling, Danvers, MA), 1:200 anti-Sox2 (DSHB, Iowa, IW) or 1:150 anti-GFAP (Cell signaling, Danvers, MA) antibodies in antibody dilution buffer (PBS, 1% BSA, 0.3% Triton X-100) buffer for overnight at 4 °C. Subsequently, the dishes were washed with PBS four times and incubated with corresponding fluorochrome-conjugated secondary antibodies in antibody dilution buffer (1:100) for 30 min. Mounting medium with DAPI (Vector) was added to the dishes which were then imaged using confocal (LSM880) or epifluorescence microscopy (Nikon). GFAP positive cells were counted under a 40X amplification field either manually or in a Keyence (Itasca, IL) microscopy system.

### Reverse transcription qualitative polymerase chain reaction

Total RNA from cells in 24-well cell culture plates was isolated using TRIZOL (Life Technologies, Grand Island, NY, USA) following the manufacturer’s instructions. Total RNA (500 ng) was converted to cDNA by Superscript III Direct cDNA Synthesis System (Life Technologies). Primers were designed to span exon-exon junction in order to preclude the amplification of genomic DNA; primers for Human *GFAP*: forward, 5’- AGGGGGCAAAAGCACCAAAGA-3’, reverse 5’CTGGGAAAATGA CGCAGTCCAG-3’; for *GDNF*: forward, 5’-CCCGCCGCAAATATGCCAGA-3’, reverse, 5’-GTTCCTCCTTGGTTTCATAGCCC-3’; for *BDNF*, forward, 5’-GCCATCCCAAGGTCTAGGTG-3’, reverse, 5’- GTGGGATGGTGGGCATAAGT-3’; for *GAPDH*; forward 5’-AACAGC GACACCC ACTCC TC-3’, reverse 5’-AGCCAAATTCGTTGTCATACCAGG-3’. The target amplicon sizes were 274, 208, 261 and 100 bp for *GDNF, BDNF*, *GFAP* and *GAPDH*, respectively. QPCR was done with ViiA Real-time PCR system (ThemoFisher). Data was analyzed with ViiA 7 software with *GAPDH* as internal control. Gene expression was normalized to vehicle control.

### Capillary Western Assays

After 30 days of differentiation, Astrocytes were treated with vehicle, DHA, EPA and SA as described above for three days and then washed twice with ice-cold PBS prior to lysis. For the standard method, cells were incubated on ice for 15 min in RIPA lysis buffer (Cell Signaling, Danvers, MA) supplemented with Complete Protease Inhibitor Cocktail (Roche) and phosphatase inhibitor (Thermo Scientific). Lysates were scraped into tubes and clarified by centrifugation at 15000 x g for 10 min at 4 °C. Total protein concentrations of the supernatants were determined by Pierce BCA Protein Assay (Thermo Scientific).

Reagents and equipment were purchased from Protein-Simple (San Jose, CA) unless stated otherwise. All experiments followed the manufacturer’s instructions. 1 µl 5X Fluorescent Master Mix was added to each sample (4 µl). Samples were incubated at 95 °C for 5 min to denature. Three microliters of each sample were loaded into the top-row wells of plates that were designed to separate proteins of 12-230 kDa. Primary antibodies were diluted in Antibody Diluent as follows: anti-BDNF (1:50), anti-GDNF (1:50) (Santa Cruz Biotechnology, Dallas, TX), anti-GPADH (1:10000) (Protein Tech, Rosemont, IL), anti-CREB (1:50), anti-pCREB (1:50) (Invitrogen, Madison, WI) and anti-GFAP (1:50) were diluted. Secondary antibodies, Anti-Mouse Secondary HRP Conjugate and Anti-Rabbit Secondary HRP Conjugate (Protein Simple). Peak area calculations were conducted with companying Compass software as indicated by manufacture (Protein Simple).

### cAMP measurement in living cells

Astrocytes differentiated for 30 days were cultured on glass bottom dishes coated with Geltrex and infected with cADDIS BacMam virus encoding the green upward cAMP sensor at 1.00 × 10^9^ VG/mL (Montana molecular, Bozeman, MT) with sodium butyrate at a final concentration of 1mM for 24 h, then replaced by astrocyte medium. Live cell images were acquired under a 20× objective on a Zeiss 880 microscope. The acute effect of n-3 PUFA on cAMP formation was evaluated for acute (20 min) DHA treatment. Every 5 min, the average fluorescent density from each cell selected from the visual field was collected. The effect of n-3 PUFAs treatment on the potential of Gαs-stimulated cAMP production was determined as previous reported [31]. Astrocytes derived from the SSRI-sensitive patient were treated for 3 days, The average fluorescent density of selected cells (5-12 cell in each dish) were first recorded, then cells were challenged with forskolin (2μM), and recording was continued. Fluorescence density was normalized to baseline fluorescence for each experiment.

### Statistical analysis

Statistical analyses were performed using the Prism 5 statistical package (GraphPad Software, Inc., La Jolla, CA). All data were analyzed by Student’s t-test or one-way analysis of variance (ANOVA) with a Tukey’s post hoc test. Data were expressed as Mean ± SD. A P value of<0.05 was considered statistically significant.

## Results

### n-3 PUFAs facilitate astrocyte differentiation from iNSCs

Decrease in astrocyte count and GFAP expression are seen at many brain regions in MDD post-mortem studies [32–35] and animal models of depression [8]. A recent reprot suggested that antidepressants could induce recovery of astrocytes in MDD, which could be a potential mechanism for their anti-depression effect [8]. To test whether n-3 PUFAs affect astrocyte formation, we observed the effect of n-3 PUFAs on astrocyte differentiation from iNSCs derived from SSRI-sensitive or -resistant MDD subjects. First, patient-derived iNSCs were confirmed with Sox2 and Pax6 co-immunostaining (Supplemental figure 1 A). The iNSCs were cultured with astrocyte differentiation medium that contained DHA, EPA, SA or vehicle for 5 days, after which cells underwent immunostaining with GFAP antibody. Note that GFAP staining is weak in some cells after 5 day differentiation, but positive staining can be identified compared to cells without expression of GFAP in the same field (Figure 1B). Cells with positive GFAP staining were counted using confocal microscopy with 40x magnification. Four fields in each dish were selected, randomly, for cell counting. Data indicated that the percentage of GFAP positive cells in each 40X field, DHA or EPA treatment significantly increased the percentage of GFAP positive cells compared to vehicle or SA treatment (Figure 1B). *GFAP* mRNA expression in cells treated with DHA or EPA was almost two times higher than in vehicle control cells (Figure 2 A). Furthermore, GFAP protein expression also increased in DHA or EPA treated cells (Figure 2 B and C). We observed that differences in astrocyte differentiation between cells from SSRI-sensitive and SSRI-resistant patients were not associated with n-3 PUFA treatment (Figure 1 and 2).

**Figure 1.**
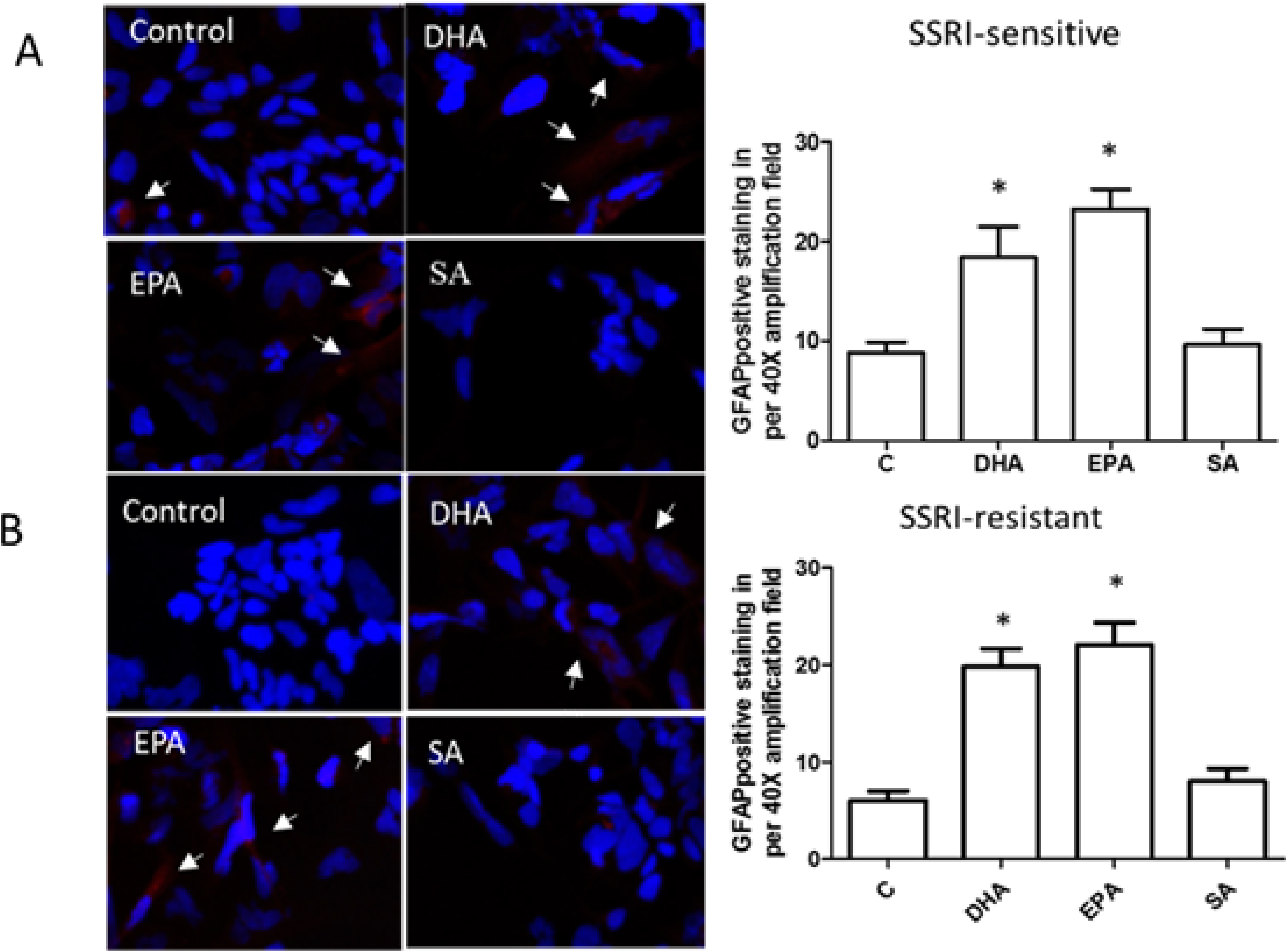
DHA and EPA treatment increased astrocyte differentiation from patient-derived iNSC. iNSC were cultured with astrocyte medium in presence of vehicle control (ethanol), DHA (docosahexaenoic acid), EPA(eicosapentaenoic acid) and SA (stearic acid) respectively for 5 days. Cells were fixed and immunostained with GFAP (Red). Arrows indicate positive cells. **A**. cell images and counts from SSRI-sensitive subject (N=3, cells in 16 fields were counted). **B**. cell images and counts from SSRI-resistant subject (N=3, cells in 16 fields were counted). * p<0.05, compared to vehicle control.

**Figure 2.**
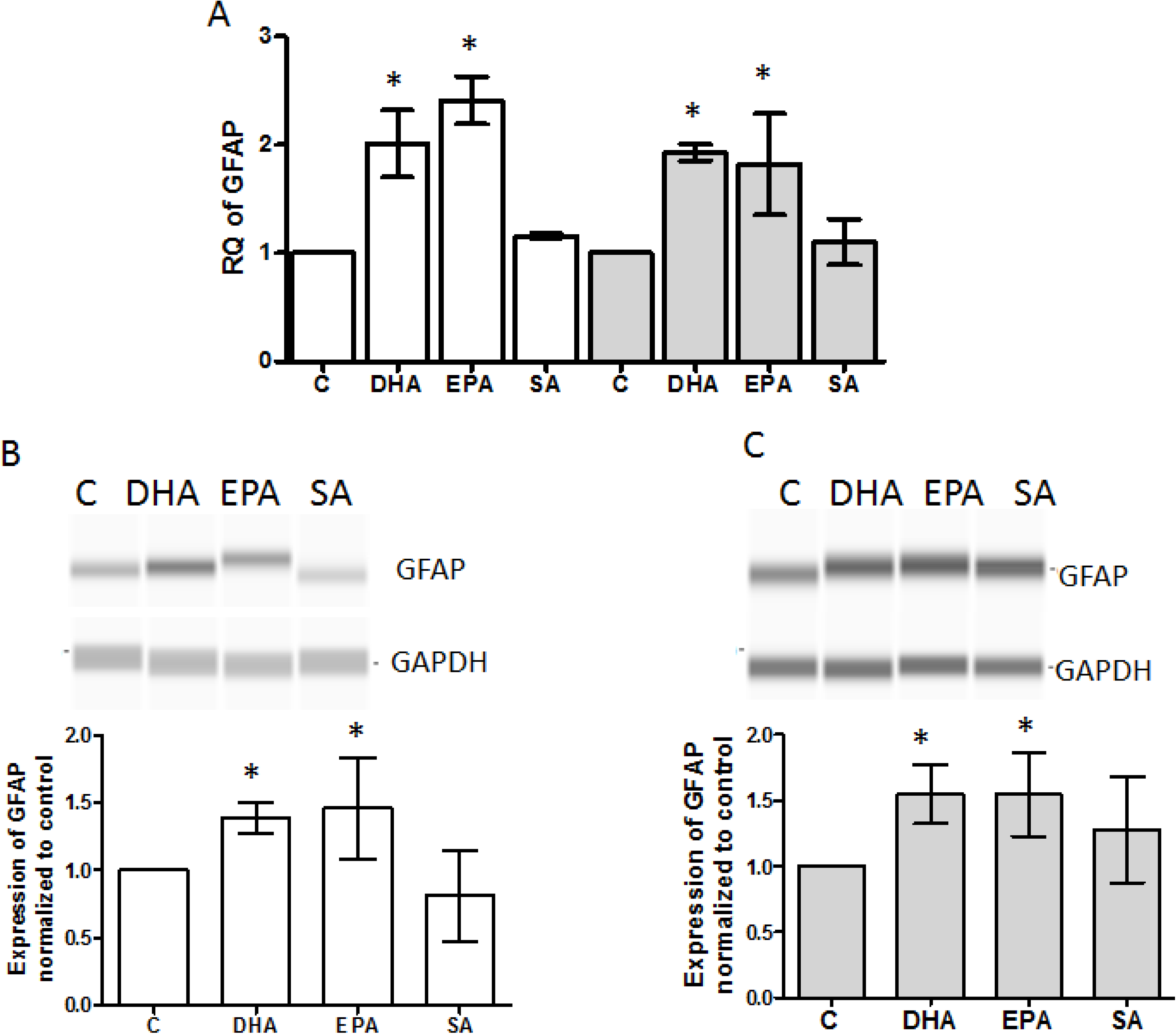
GFAP expression was increased by n-3 PUFAs treatment during astrocyte differentiation from patient-derived iNSC. iNSC were treated with DHA, EPA, SA and vehicle control for 5 days in astrocyte medium, and then tested for GFAP mRNA and protein expression. **A**, mRNA expression of GFAP was quantified using qPCR (N=3). The white bar represents data from SSRI sensitive subject and the gray bar from the SSRI resistant subject. GFAP protein in cells from SSRI-sensitive subject (**B**) and SSRI-resistant subject (**C**) was determined with immunoblot using capillary western blots (N=5) * P<0.05 compared to vehicle control.

### N-3 PUFAs increased BDNF and GDNF production in astrocytes from patient-derived iNSC

Therapeutic effects of antidepressants on MDD by modulation of astrocytic function (such as producing BDNF and GDNF) have been reported [36–39]. Though animal and clinical studies on MDD showed increased BDNF with n-3 PUFA treatment [40–42], the relevant cell types were not delineated. Here, astrocytes were generated with patient-derived iNSCs. After 30 days of differentiation in astrocyte differentiation medium, about 80-90% of cells were GFAP positive (Supplemental figure 1 B). The astrocytes were cultured with vehicle control, DHA, EPA and SA for an additional 3 days. mRNA expression for BDNF or GDNF were tested. DHA and EPA significantly increased the mRNA of BDNF and GDNF in astrocytes derived from SSRI- sensitive and resistant patients, SA was without effect (Figure 3 A and B). The results from capillary western assay further demonstrated that DHA and EPA elevated BDNF and GDNF protein in the astrocytes from both SSRI-sensitive and SSRI-resistant subjects (Figure 3 C). This increase in BDNF and GDNF induced by DHA and EPA required sustained n-3 PUFA treatment and was not evident after a 24 h treatment. (Supplemental figure 2).

**Figure 3.**
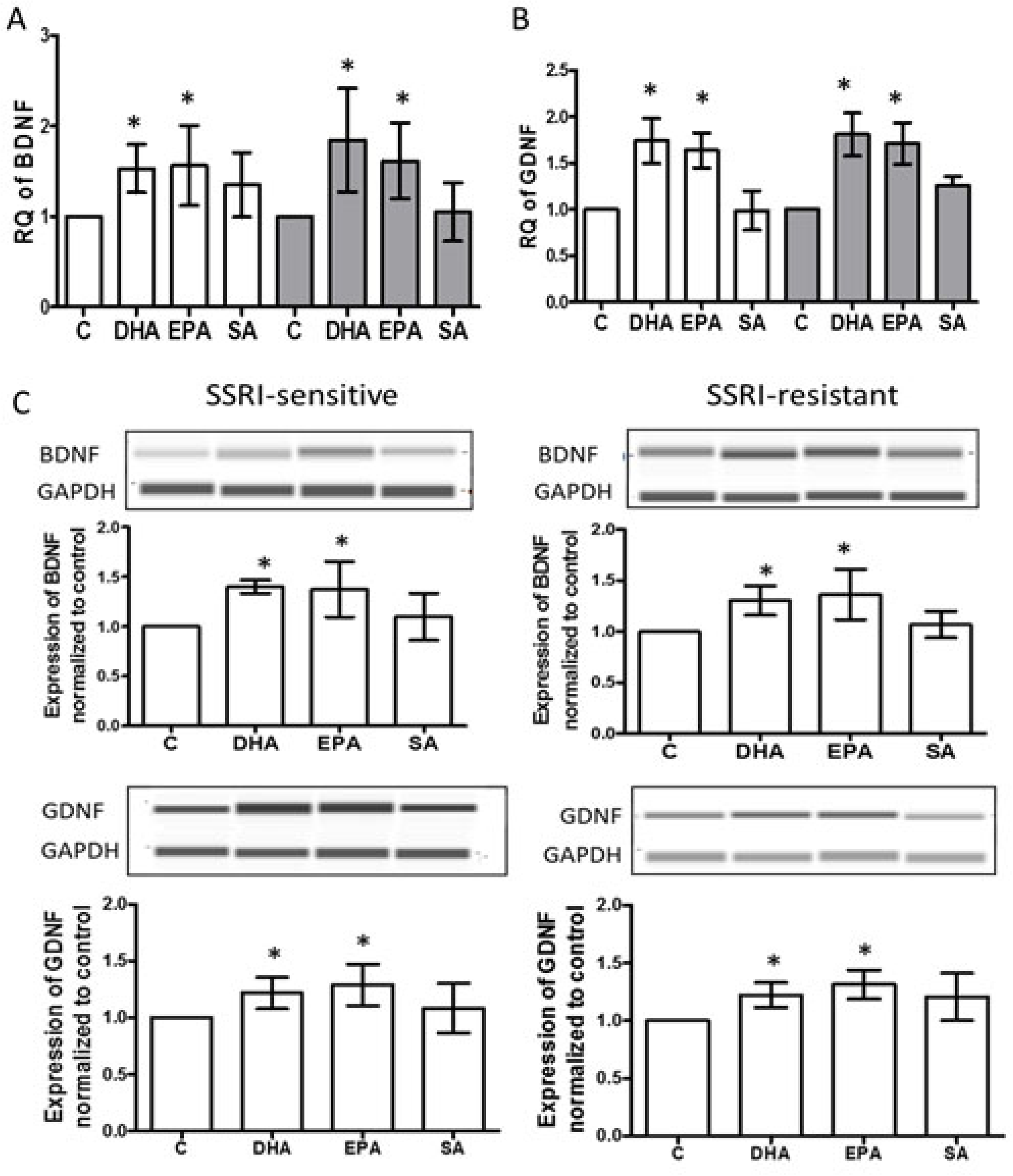
N-3 PUFAs elevated BDNF and GDNF production in astrocytes from patient-derived iNSC. After treatment, as indicated, for 3 days. BDNF (A) and GDNF (B) mRNA expression were measured with qPCR. White bar represents astrocytes derived from SSRI- sensitive patient and gray bar, from SSRI-resistant patient (N=6). * p<0.05, compared to vehicle control. C, Astrocytes were treated with vehicle control, DHA, EPA, and SA, as indicated, for 3 days. Cell lysates were subjected to capillary western assay. Data were processed with software provided by the manufacturer (ProteinSimple). The expression of BDNF or GDNF (represented by peak area) was normalized to vehicle control. Each experiment was repeated at least 4 times *p<0.05, compared to vehicle control

### PUFAs increased phosphorylated CREB in astrocytes in a cAMP independent manner

CREB is an important regulator for BDNF and GDNF expression [43–45]. Chronic antidepressant treatment enhances CREB activation, suggesting CREB as an important mediator for antidepressant treatment [24]. CREB must be phosphorylated to pCREB in order to transcribe CREB-regulated genes, including BDNF and GDNF [46].To examine whether n-3 PUFAs alter the activity of CREB in patient-derived astrocytes, CREB and pCREB were measured using a capillary western assay in 30-day differentiated astrocytes. After 3 days treatment with DHA, EPA, SA and vehicle control, CREB expression did not show significant alteration, but pCREB was increased in both astrocytes derived from SSRI-sensitive and -resistant subjects (figure 4), suggesting that DHA and EPA elevated CREB activity. cAMP activates CREB through protein kinase A (PKA). Antidepressants, like escitalopram, evoke a sustained increase in cAMP [47]. The patient-derived astrocytes were treated with RP-cAMP (150 uM) +vehicle, DHA, EPA or SA, respectively for 3 days. RP-cAMP, a potent and specific competitive inhibitor of the activation of PKA by cAMP, did not diminish the increase of pCREB induced by DHA or EPA (Figure 4), suggesting that cAMP cascades may not be involved in activation of CREB by n-3 PUFAs. With RP-cAMP treatment, DHA and EPA still elevated the BDNF and GDNF production (Supplemental figure 3). Antidepressants including ketamine, translocate Gα_s_ from lipid rafts, which increases association of Gα_s_ with adenylyl cyclase, increasing cellular cAMP production [31, 47]. To test if n-3 PUFAs have similar effect, we measured the cAMP with a fluorescent sensor in astrocytes from SSRI-sensitive and -resistant subjects. After three days of treatment, no difference was seen among baselines of each group. However, DHA and EPA increased Gαs-activated cAMP production stimulated by forskolin, while SA did not show the effect in SSRI-sensitive cells (Supplemental figure 4). Similar to antidepressants, after 20 min treatment with n-3 PUFAs, no change in cAMP was seen (Supplemental figure 5)

**Figure 4.**
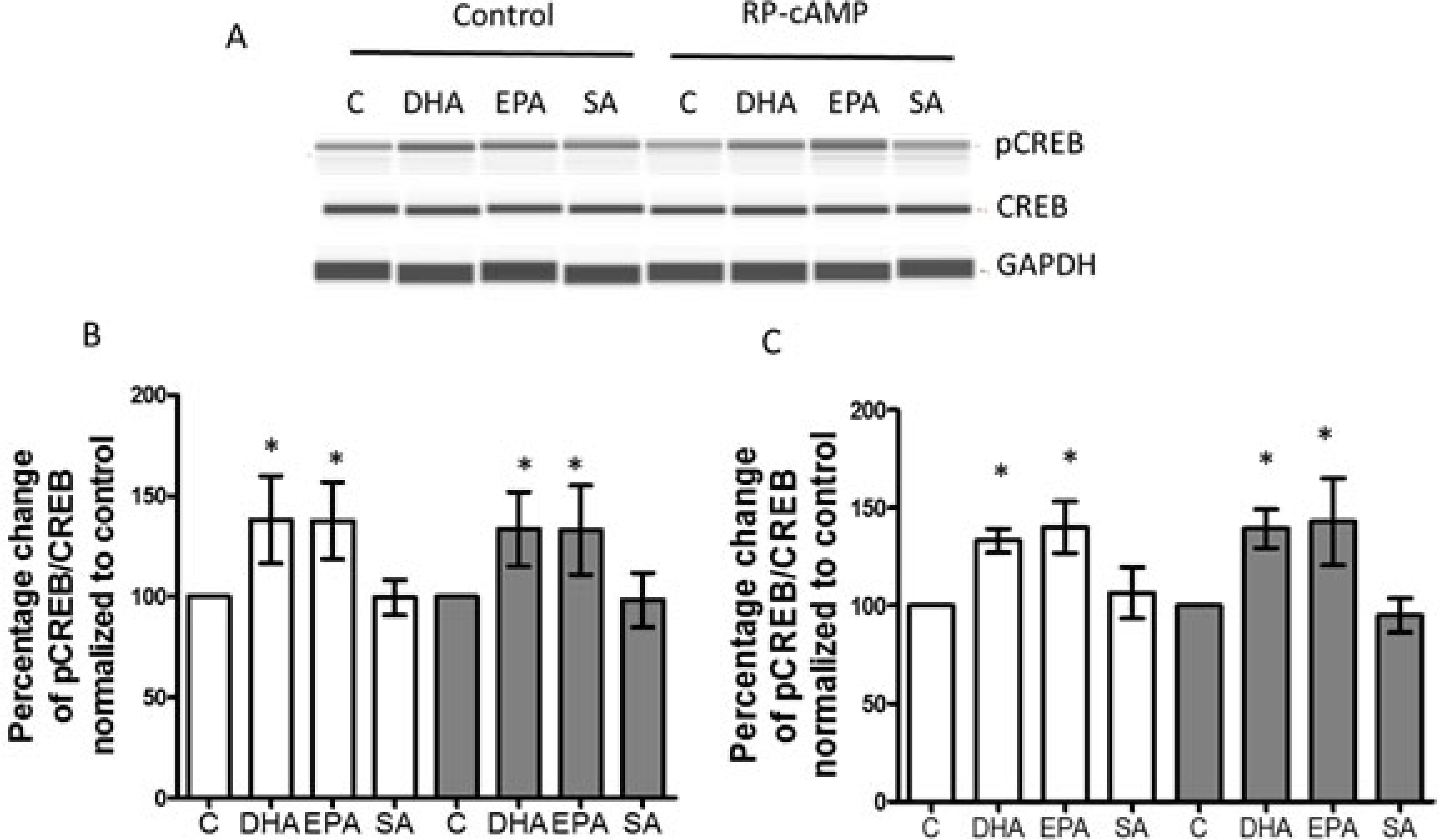
CREB was activated by DHA or EPA treatment, and this was cAMP independent. **A**, Astrocytes from patient-derived iNSC were treated with vehicle control, DHA, EPA, and SA for 3 days. Cell lysates were used for capillary western blots and a representative image is shown. RP-cAMPS, a cAMP antagonist, was used at 150 µM. **B**, Quantified data from capillary western assays. Cells were treated with vehicle control, DHA, EPA and SA as indicated. **C**, Quantified data for phosphorylated CREB. Cells were treated with vehicle control, DHA, EPA and SA ± RP-cAMP for 3 days. White bar, SSRI sensitive subject, Gray bar, SSRI resistant subject (N=4). The ratio of pCREB with CREB was normalized to vehicle control. *p<0.05, compared to vehicle control.

**Figure 5.**
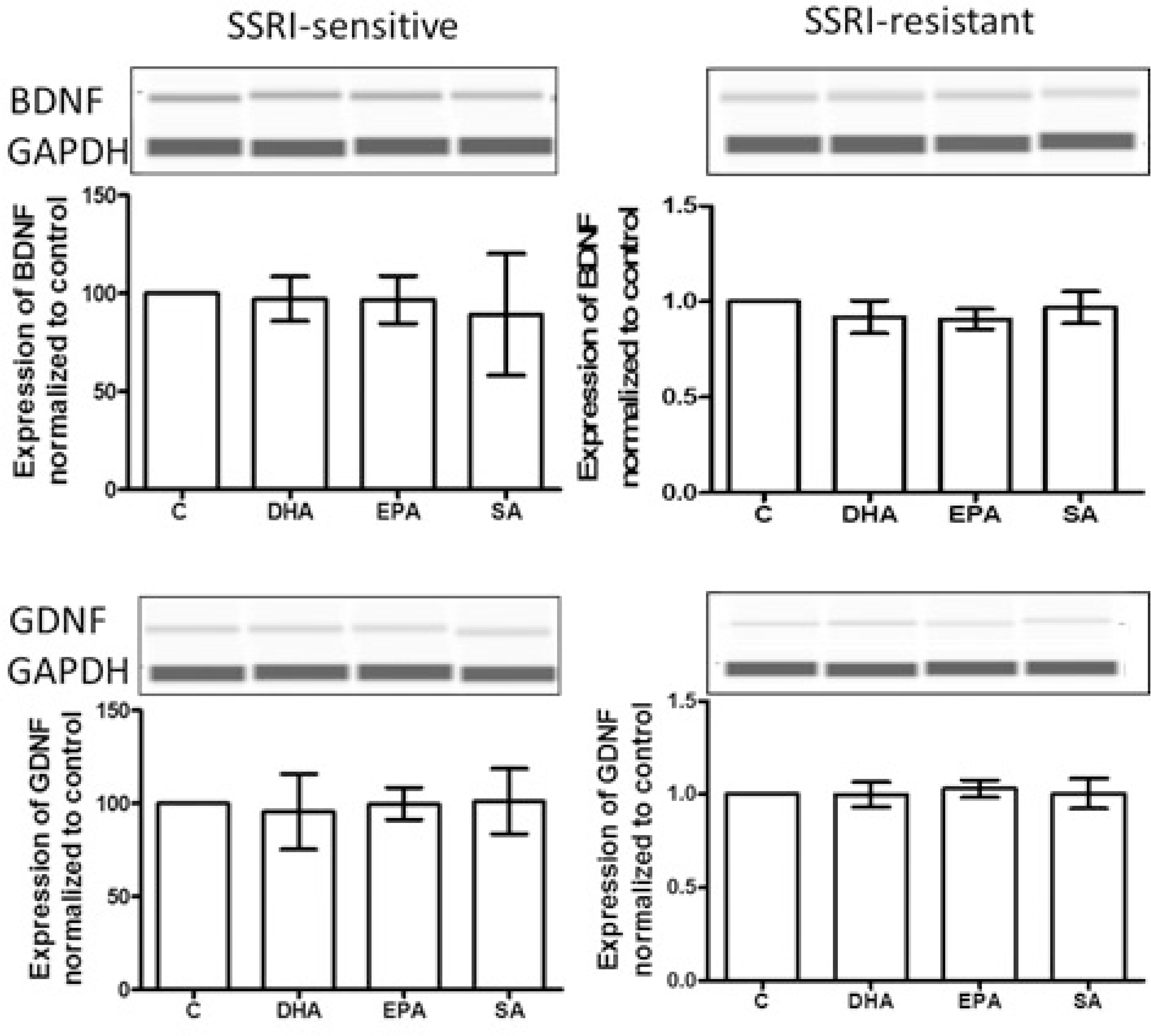
CREB inhibitor attenuated the BDNF and GDNF increase induced by n-3 PUFAs in astrocytes. Astrocytes were treated with vehicle, DHA, EPA and SA for 2 days, then the CREB inhibitor, 666-15 (10 μM), was applied for 1 day. The 3-day treated cells were collected and analyzed in capillary western blots. Expression of BDNF and GDNF showed no significant alteration among different treatments. Each experiment was repeated 4 times.

### 4. Increased expression of BDNF or GDNF by n-3 PUFAs in patient-derived astrocytes is mediated by CREB

CREB/pCREB and BDNF appear to be diminished in animal models and in patients suffering from depression [37]. In a CREB-deficient mouse, the upregulation of BDNF induced by antidepressant was abrogated [48]. To explore whether CREB mediates the elevated BDNF and GDNF production induced by n-3 PUFAs, astrocytes were treated with vehicle, DHA, EPA and SA for two days, then 666-15 (10 μM), a specific inhibitor of CREB [49], was added and cells were cultured for an additional 24 hours. The cellular lysate was subjected to capillary western assay. BDNF or GDNF production induced by DHA or EPA was attenuated by 666-15 in astrocytes that were derived from SSRI-sensitive or SSRI-resistant subjects. (Figure 5).

## Discussion

N-3 PUFAs play fundamental roles in brain structure and function. Accumulating evidence has implicated deficiency of n-3 PUFA (EPA and DHA) in the etiology of depression and n-3 PUFA in the treatment of MDD [13, 15]. Multiple mechanisms might underlie the protective/therapeutic effect of n-3 PUFAs on MDD [13–17]. Decrease in astrocyte count and GFAP expression are often seen in MDD post-mortem studies [32, 33] [34, 35] [50]. Astrocyte reduction was also observed in a variety of animal models of depression [8]. Antidepressants, including SSRIs, could increase/restore the astrocyte counts or GFAP staining, which positively correlated with attenuated “depression-indicators” in rodents [8]. These suggested a mechanism for relief of depression by restoring astrocyte function. Some reports have indicated that n-3 PUFA treatment could impact the neuronal differentiation in primary cultured animal NSCs [51]. A recent report indicated that DHA could enhance astrocyte-like morphologic formation in rat astrocytes [52]. Here, taking the advantages of iPSC technology, the effect of n-3 PUFAs on astrocyte differentiation from patient-derived iNSCs was tested. These data suggested that n-3 PUFAs could help restore the astrocyte population in MDD since n-3 PUFAs facilitated astrocyte differentiation from patient-derived iNSC, as measured by an increase in GFAP expression and GFAP positive cell counts compared to cells without n-3 PUFAs treatment. Thus, gliogenesis may be relevant to therapeutic effects of n-3 PUFAs.

Human studies and animal models support relevance of neurotrophins to depression, proposing that MDD is associated with decreased expression and/or function of BDNF and GDNF, which can be alleviated with antidepressant therapy [53, 54] (though there are some inconsistencies [38, 55]). Furthermore, overexpression of BDNF in astrocytes leads to antidepressant-like activity in mice [12]. Ketamine elevated BDNF in cultured mouse astrocyte, and did this along a more rapid time course than monoamine-centric antidepressants [31]. N-3 PUFAs appeared to have similar effect on neurotrophin production as antidepressants. DHA and EPA elevated BDNF and GDNF expression after three day treatment, but the saturated fatty acid, SA, did not. This is similar to the treatment time required for antidepressants to translocate Gsα from lipid rafts in C6 cells [18].

It is worthy to mention that many (but not all) of these studies show greater efficacy of EPA than DHA, while our studies suggest equal potency and efficacy for the two compounds [14, 52, 56–58]. Note, however, that 1) the EPA- or DHA-“enriched” preparations used in clinical studies are 80:20 mixtures of the two n-3 PUFA species and 2) orally-ingested preparations are metabolized, something that does not occur in cell-culture medium

Several reports have shown that n-3 PUFAs could increase CREB expression and pCREB in depression animal models [59]. In agreement with these reports, DHA and EPA increased pCREB in patient-derived astrocytes, except that CREB expression was not altered by n-3 PUFAs in this study. The difference may result from using a different experimental model (patient-derived astrocytes versus animal brain). CREB can be activated by cAMP-PKA signaling pathway. The data from us and others have shown that chronic antidepressant treatment increased adenylyl cyclase (AC) activity and cAMP in cells [31, 47, 60]. Phosphodiesterase 4 (a cAMP-specific isoform) inhibitors show antidepressant effects in both patient and rodent models [61–64]. Activation of CREB via the cAMP-PKA signaling pathway is thought to mediate the therapeutic effects of antidepressants [24]. However, results from this study suggest that cAMP-PKA does not participate in n-3 PUFAs-activated CREB in patient-derived astrocytes (figure 4). Recently, a study reported an increase of cAMP subsequent to brief EPA treatment in epithelial cells [65]. We did not observe this in the human astrocyte preparation, so a cAMP increase induced by n-3 PUFAs might be cell type-dependent. Notably, data have indicated that antidepressants, including ketamine, could increase cAMP by regulating interaction of Gαs with adenylyl cyclase [31, 47]. N-3 PUFAS have similar effect; after 3-day treatment with n-3 PUFAs, Gαs-stimulated cAMP production was increased.

Consensus-binding sequences for CREB were identified in the promoter sequence of either *BDNF* or *GDNF* genes [66]. Several studies suggested that elevation of expression of BDNF or GDNF induced by antidepressants was dependent on CREB activity [37, 48], however, this has not been documented for n-3 PUFAs. In this study, the cell permeable CREB specific inhibitor[49], indicated that pCREB mediated the elevation of BDNF and GDNF induced by DHA or EPA in the astrocytes, even as the process was cAMP-independent.

Note that there are some limits to data interpretation in this study. Firstly, astrocytes were derived from stem cells obtained from two subjects, one SSRI-sensitive and the other, SSRI-resistant. Both appear to have the same response to n-3 PUFA treatment in astrocyte differentiation, production of BDNF and GDNF, and activation of pCREB. It is possible that these subjects would have both responded positively to n-3 PUFA, but this cannot be established. In addition, though astrocytes derived from iPSCs provide many advantages for *in vitro* depression studies, the induced astrocytes are generally considered developmentally immature. These cells cannot predict the behavior of the subjects from whom they were derived.

In summary, this study suggests that DHA and EPA could facilitate astrocyte differentiation from iNSCs and like antidepressants, increase BDNF and GDNF production in those cells. While pCREB mediates this effect, it is cAMP-independent. The study further adds to a growing body of work suggesting that iPSC technology is a useful tool to probe the cellular basis of MDD and antidepressant response. This study also further establishes a role for astrocytes in depression and antidepressant therapy.

## Acknowledgements

This research is supported by NIH R01AT009169 (MMR, JZY) and VA Merit Award BX00149 (MMR). Patient cell line collection and derivation were supported by NIH P50MH106933 and NIH R01AT009144 (RHP, SDS, JW).

## Figure legend for supplemental data

**Supplemental Figure 1.**
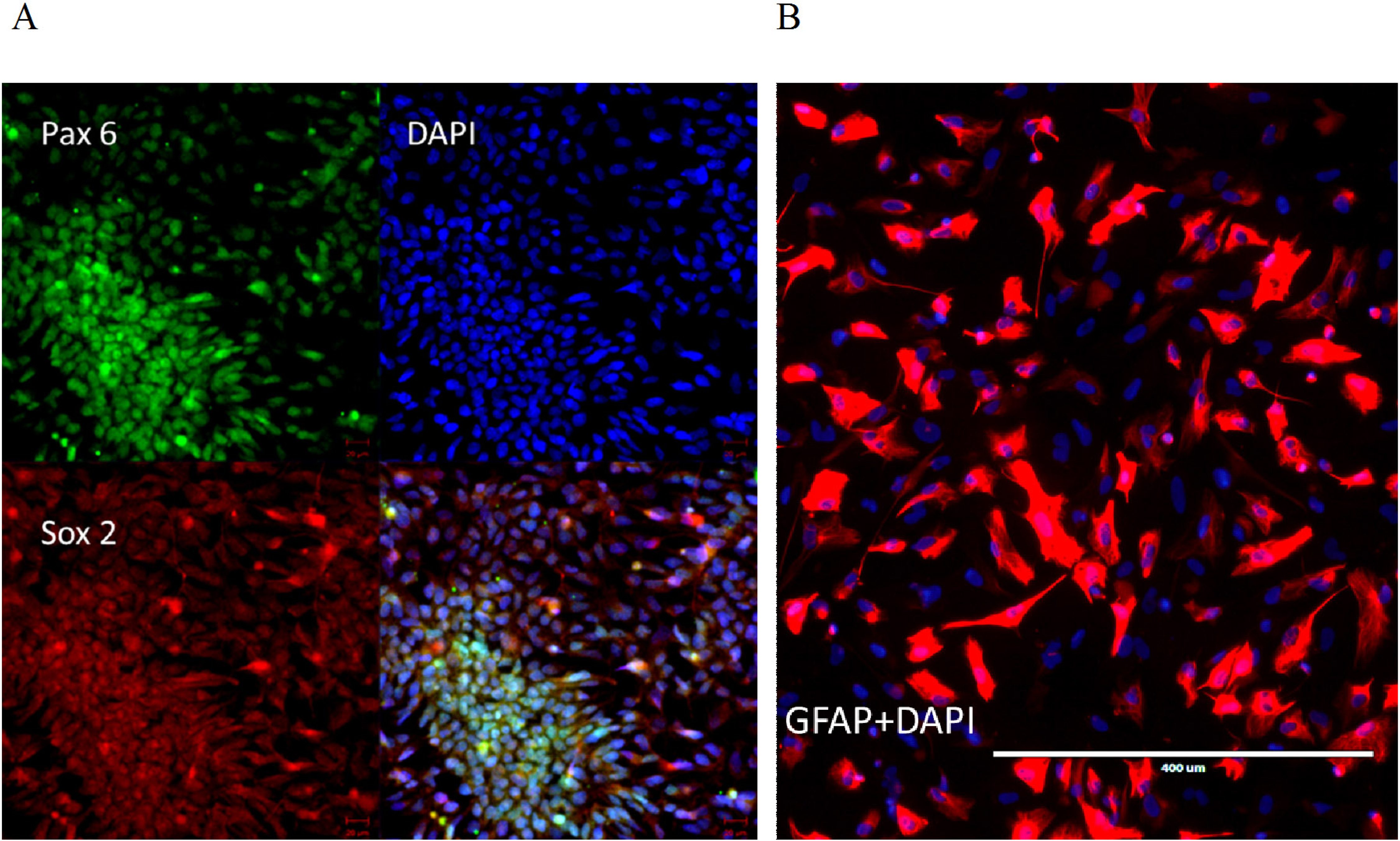
Characterization of patient-derived iNSC and astrocytes. **A.** representative images for PAX6 (green) and SOX2 (red) immunostaining of iNSC. Nuclei are labeled with DAPI (blue). Image was taken with LSM 880 confocal microscopy using 20x magnification. **B**. GFAP immunostaining for astrocytes. Red indicates GFAP positive, blue is DAPI staining for nuclei. Image was obtained with epifluorescence microscopy.

**Supplemental Figure 2.**
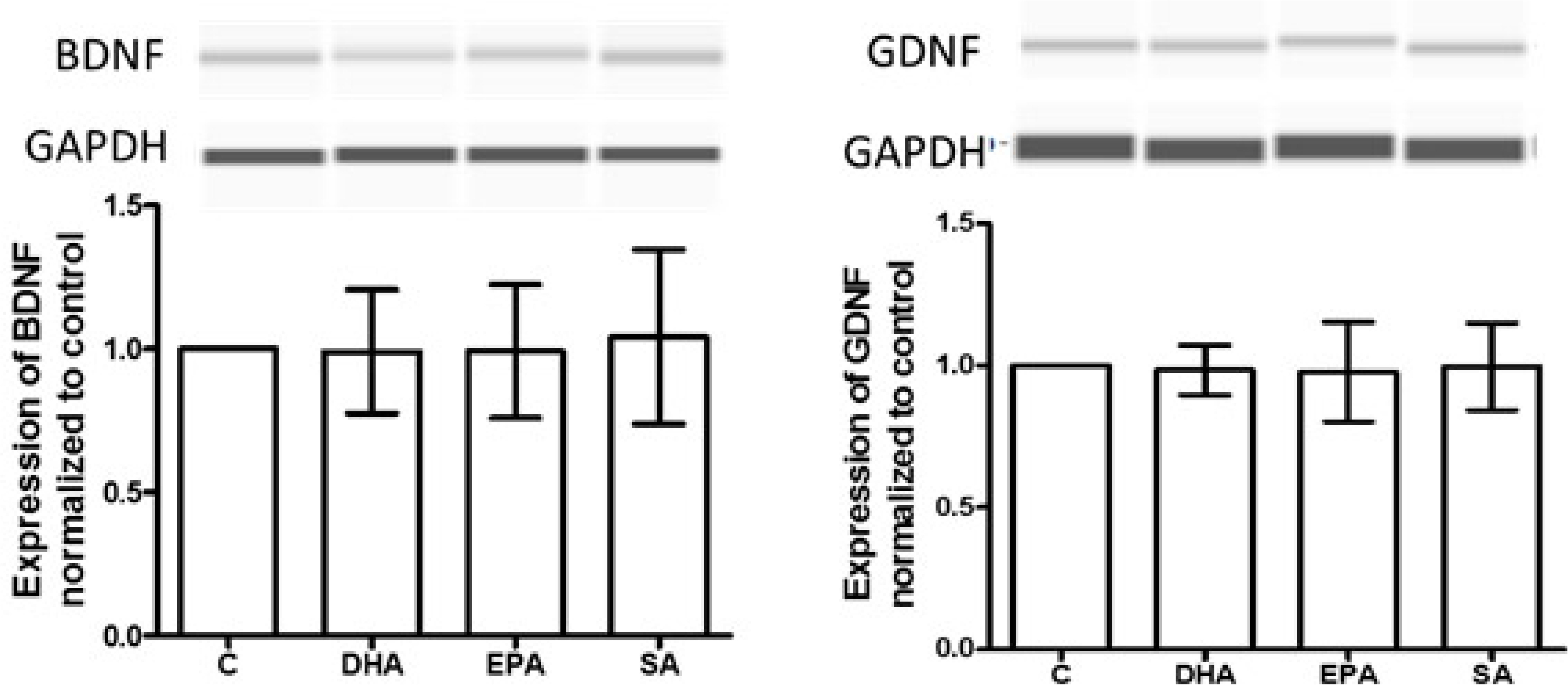
Brief (24h) n-3 PUFAs treatment did not alter BDNF and GDNF production. Astrocytes from SSRI-sensitive subject-derived iNSC were treated with n-3 PUFAs for 1 day, and cells were lysed and processed for capillary western assay. DHA, EPA and SA did not alter BDNF and GDNF production, compared to vehicle control (n=3).

**Supplemental Figure 3.**
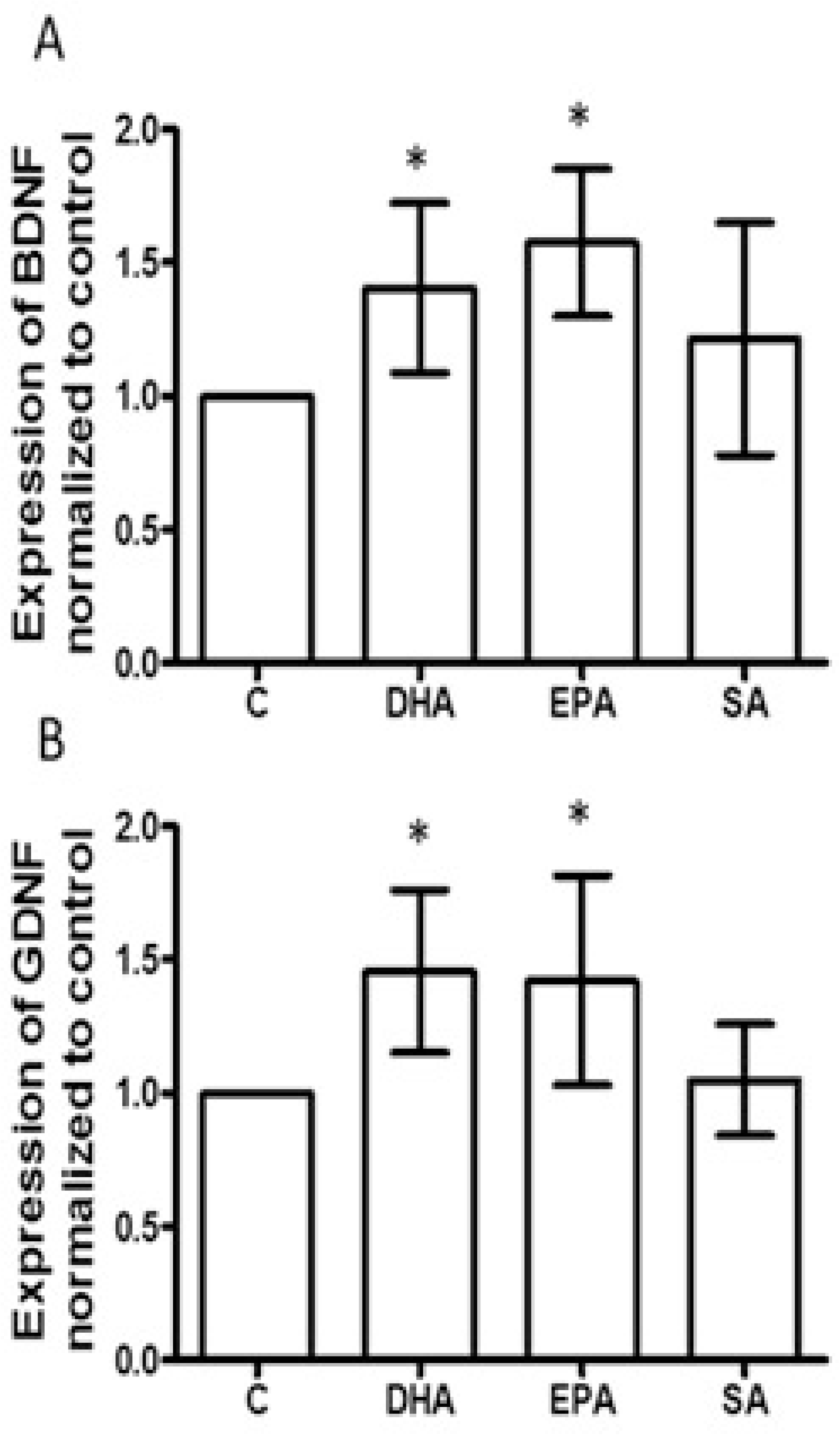
RP-cAMPS did not affect BDNF or GDNF expression induced by n-3 PUFAs in astrocytes. Astrocytes from iNSC derived from an SSRI-responsive patient treated with reagents as indicated in figure 4 and RP-cAMP for 3 days. Cell lysates were used for capillary western assay. **A**, Quantified data for BDNF in astrocytes derived from an SSRI- responsive subject (N=4). **B,** Quantified data for GDNF in cells from SSRI-resistant subject (N=4). *p<0.05, compared to vehicle control,

**Supplemental Figure 4.**
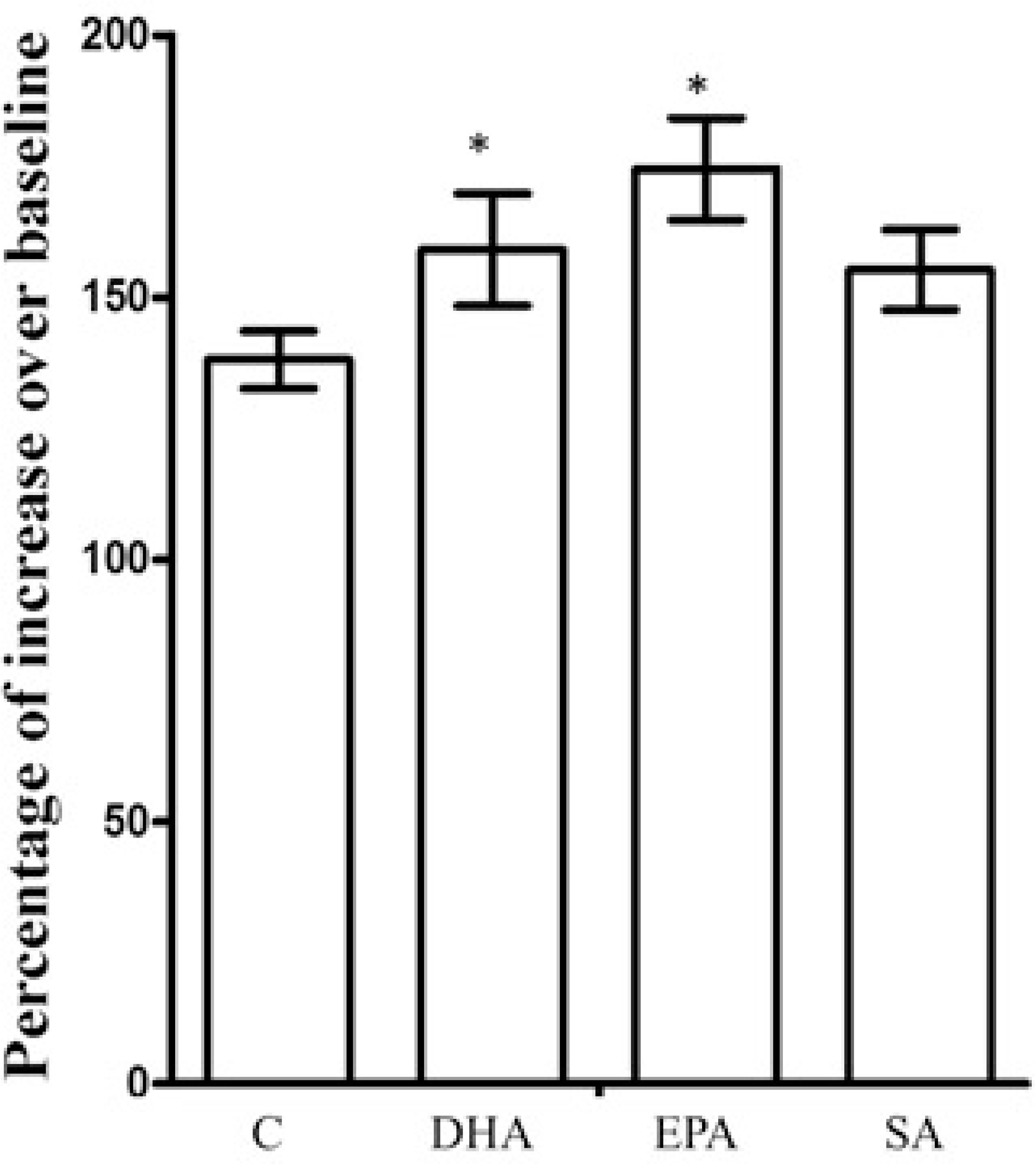
Three day N-3 PUFAs treatment increased cAMP potential with forskolin challenge, After 3-day treatment with vehicle (Control), DHA, EPA and SA, Astrocytes were infected with (1.09 × 10^9^ VG/mL) cADDIS virus. The basal florescent density of the selected cells were recorded first, then cells were stimulated with forskolin (2 μM), and immediately the fluorescent density of cell was determined. cAMP alteration in cell was represented by percentage of increase over baseline. *p<0.05. Compared to vehicle control, n=4.

**Supplemental Figure 5.**
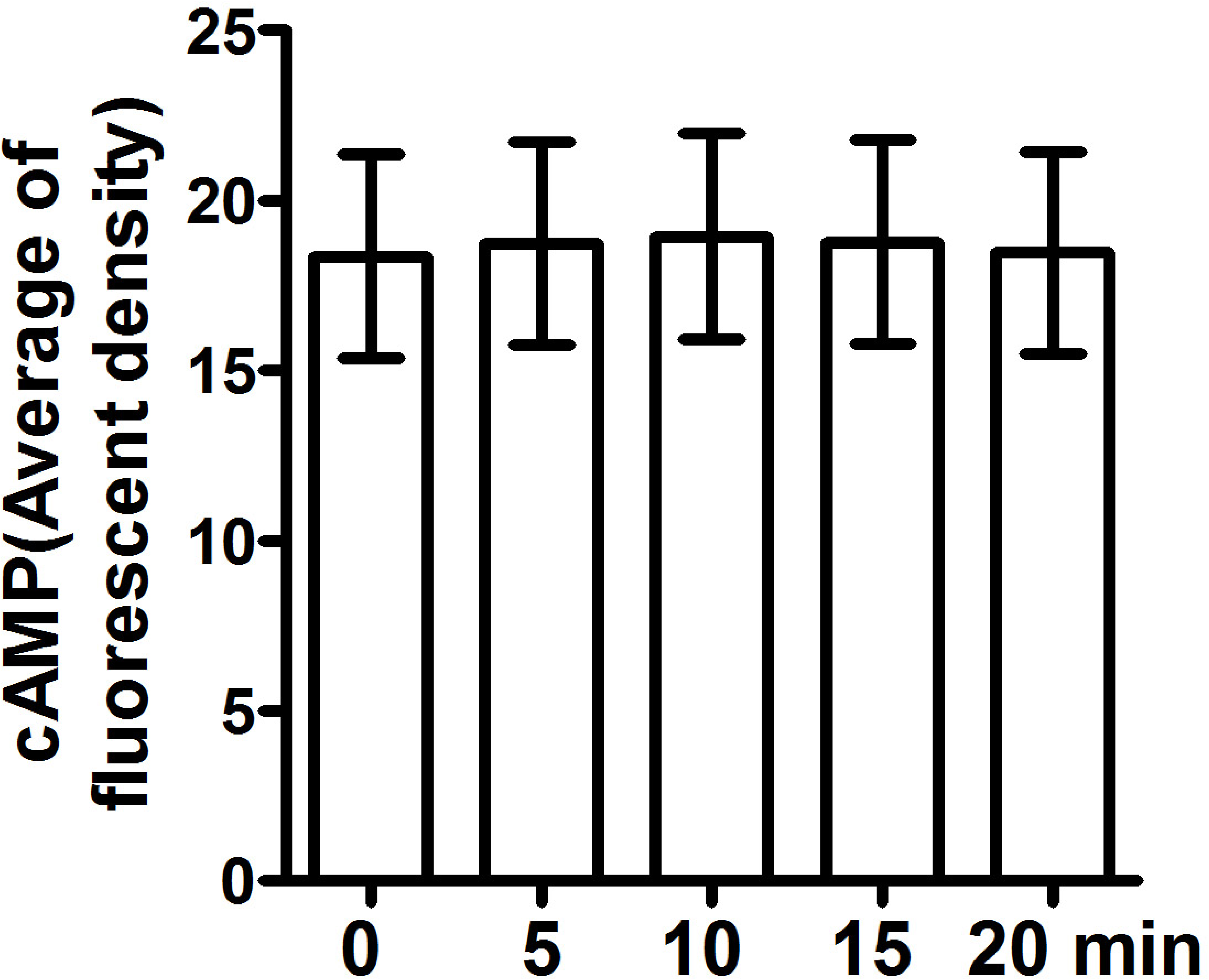
DHA did not alter astrocyte cAMP production during 20 min treatment. Astrocytes were infected with (1.09 × 10^9^ VG/mL) cADDIS virus. The florescent density of the selected cells were recorded with confocal microscopy (LSM-880) every 5 min. The average fluorescent density at each time point measured (0, 5, 10, 15, 20 min) was roughly equal. (n=3)

